# Effect of composting and storage on the microbiome and resistome of cattle manure from a commercial dairy farm in Poland

**DOI:** 10.1101/2023.12.04.569838

**Authors:** Magdalena Zalewska, Aleksandra Błażejewska, Mateusz Szadziul, Karol Ciuchciński, Magdalena Popowska

## Abstract

Manure from food-producing animals, rich in antibiotic-resistant bacteria and antibiotic resistance genes (ARGs), poses significant environmental and healthcare risks. Despite global efforts, most manure is not adequately processed before use on fields, escalating the spread of antimicrobial resistance. This study examined how different cattle manure treatments, including composting and storage, affect its microbiome and resistome. The changes occurring in the microbiome and resistome of the treated manure samples were compared with those of raw samples by high-throughput qPCR for ARGs tracking and sequencing of the V3-V4 variable region of 16S rRNA gene to indicate bacterial community composition. We identified 203 ARGs and mobile genetic elements (MGEs) in raw manure. Post-treatment reduced these to 76 in composted and 51 in stored samples. Notably, beta-lactam, cross-resistance to macrolides, lincosamides and streptogramin B (MLSB), and vancomycin-resistance genes decreased, while genes linked to MGEs, integrons, and sulfonamide resistance increased after composting. Overall, total resistance gene abundance significantly dropped with both treatments. During composting, the relative abundance of genes was lower midway than at the end. Moreover, higher biodiversity was observed in samples after composting than storage. Our current research shows that both composting and storage effectively reduce ARGs in cattle manure. However, it’s challenging to determine which method is superior, as different groups of resistance genes react differently to each treatment, even though a notable overall reduction in ARGs is observed.

## 1 Introduction

The increased demand for food of animal origin has resulted in growing numbers of food-producing animals producing large quantities of manure; indeed, a single dairy cow can produce 54 kg of wet manure per day (Girotto and Cossu, 2017). Such production has raised environmental concerns which are driving food animal producers to identify economically and environmentally-safe solutions for manure disposal.

Cattle manure management is especially important for Europe, because approximately 21% of global cattle milk production takes place in the European Union (EU), with almost half of this EU production being concentrated in Germany, France, and Poland (FAO, 2020). Although, in Europe, regulations limiting antimicrobial use in animal production are restrictive, and antibiotics are applied only to treat bacterial infections, in other regions, antibiotics are still used as growth promoters (Dibner and Richards, 2005; Centner, 2016; Hu and Cowling, 2020). Even though antibiotics are used for medicinal purposes, many cases of antibiotic overuse or misuse have been reported in the animal production sector, and this has been associated with the emergence of antibiotic resistance in bacteria. After ingestion, only a small amount of the consumed antibiotics is absorbed or metabolized from the animal intestine, with about 75% of the applied antibiotic being excreted into the feces or urine: the exact percentage depends on the antimicrobial class (Sarmah et al., 2006; Amarakoon et al., 2016; Spielmeyer, 2018; Filippitzi et al., 2019). Limitation of antibiotic use seems to be a proper way to reduce the severity of antibiotic resistance occurrence (Kaur Sodhi and Singh, 2022).

Due to the breeding programs applied in the dairy industry, modern dairy cows are now highly productive; however, they are also prone to a range of infections, particularly those related to the mammary gland (Zalewska et al., 2020). In such cases, bovine mastitis, dry cow therapy, and udder health disorders in dairy cattle are often treated using various antibiotics, including cephalosporins, penicillins, and macrolide-lincosamides (Barlow, 2011). In fact, these conditions are the most common cause of antibiotic use, followed by respiratory, reproductive, and gastrointestinal diseases (Ferroni et al., 2020). Antimicrobial prescription practices vary by country, and cattle are also often administered aminoglycosides, amphenicol, quinolones, and sulfonamides, often combined with trimethoprim (Merle et al., 2012; Diana et al., 2020; Ferroni et al., 2020; Olmos Antillón et al., 2020). In addition, dairy cattle demonstrate significantly higher mean annual antibiotic consumption than beef cattle (Merle et al., 2012; Ferroni et al., 2020).

The gastrointestinal tract of dairy cattle is inhabited by a diverse bacterial community, whose composition and function play key roles in animal wellness and production efficiency. The composition of the community varies along the gastrointestinal tract; however, the most common bacteria overall belong to the phyla Bacillota (65%), Bacterioidota (15%) and Pseudomonadota (13%) (Mao et al., 2015). Although antibiotics intended to target specifically pathogens, other non-target bacteria are also affected (Mann et al., 2021). Antibiotic treatment of cattle induces shifts in the taxonomic structure and biodiversity of the gastrointestinal microbiota (Beyi et al., 2021); however, this effect varies between the compartments of the digestive tract, with the rumen being the most heavily impacted (Thomas et al., 2011). Unfortunately, this rich microbiota may also harbor antibiotic-resistance genes (ARGs). Animal feces can therefore become a source of manure-borne antimicrobial resistance (AMR) spread, especially in more intensive livestock farming scenarios, characterized by both high amounts of bacteria and significant antimicrobial selective pressure, promoting the occurrence and enhancement of antibiotic-resistant bacteria (ARB) and ARGs (Ferroni et al., 2020; Huygens et al., 2021). AMR determinants such as ARB, ARGs, and mobile genetic elements (MGEs) can contaminate surface water, groundwater, rivers, ponds, lakes, soil, or air through several pathways: (a) directly through runoff from animal production/breeding systems, (b) via the direct application of animal manure on arable fields for crop production, (c) through irrigation water from manure storage ponds and manure treatment facilities, and (d) via air particles (McEachran et al., 2015; Nguyen et al., 2020). During therapy, the antibiotic reaches the highest possible concentration in the treated body for efficient therapy without any toxic effect on the host, but when an antibiotic is excreted into the natural environment, it reaches doses sublethal for many bacteria. In this way, the antibiotic only causes a selection pressure that stimulates the bacteria to develop resistance (Kaur Sodhi and Singh, 2022). Antibiotics have been found in almost all environments, making them the pollutant of enormous concern. It has been estimated that until 2050 ARB will constitute the main cause of mortality; thus new ways to efficiently remove them from the environment as well as new tools to combat already developed AMR are needed (Kaur Sodhi and Singh, 2022; Singh et al., 2023).

Dairy cattle can produce as much as 7.3 kg of dry mass of manure per animal per day (compared to 5.5 kg in feedlot cattle), which is equivalent to 7-8% of the body weight of the cow (Font-Palma, 2019). In 2020, manure production in dairy cattle in Poland totaled approximately 149 million kg of nitrogen content, accounting for about 7.5% of production in the EU (FAO, 2020). The easiest disposal strategy for unprocessed manure is direct land application; however, such raw manure might contain antibiotic residues, ARB and ARGs, thereby increasing the risk of AMR being spread into the environment. This poses a significant threat to both human and animal health. The persistence and leaching of antibiotics into the ground and surface waters depend on the physicochemical properties of the antimicrobial compound and the fertilized soil. These compounds can persist for extended periods, and eventually accumulate in crops (Zalewska et al., 2021). The continuous presence of low concentrations of antibiotics creates selective pressure in the environment, promoting the growth of resistant bacteria (Gullberg et al., 2011). The presence of ARGs in the environment has even greater implications compared to the presence of antibiotics itself (Ahmed et al., 2021). Consequently, extensive research on the effects of manure management practices is necessary to understand their effect on microbial communities and the spread of AMR.

The choice of manure management method can influence the microbial community of the manure and reduce the survival of pathogens (Heinonen-Tanski et al., 2006). The most common strategy involves storing the feces for a prolonged time, usually between 4 to 7.5 months; however, this requires sufficient storage capacity (Loyon, 2018). An alternative strategy is the composting of manure, which is both economically and environmentally beneficial, as it reduces solid organic waste volume while eliminating pathogens, antibiotic residues and ARGs (Dolliver et al., 2007; Tasho and Cho, 2016; Gou et al., 2018). Composting transforms organic waste into a stable, humus-like product that is easily available for soil amendment. Although composting offers numerous advantages, estimates from 2011 show that only 0.8% (10.4 million tons) of total EU livestock manure was composted. Moreover, only about 8% of the manure used in fields underwent any form of treatment (Foged et al., 2012). In the Netherlands, 75% of manure from calves was applied to agricultural land without any treatment (Berendsen et al., 2018). It is believed that the primary reason for omitting such processing is the lack of adequate storage capacity (Köninger et al., 2021).

The structure of the bacterial community in cattle manure and its resistome (pool of ARGs in the bacterial community) remain relatively unknown, with a lack of comprehensive data on the subject. Until now, little studies have compared the effects of storage and composting of manure originating from the same source; only a few have examined the effects of multiple manure treatment strategies on the same raw manure batch. The aim of the study was to determine how composting and storage of dairy cattle manure impact the ARGs and microbiome of animal feces, with the goal of limiting the potential risk of spreading ARGs in the fields and compare these both manure treatment techniques in term of better preparation manure for land application. The material was collected simultaneously from the same animals, allocating one portion of the sample for storage and another for composting. The analysis considers the type of process as the independent variable, with ARGs relative abundance and bacterial phylum relative abundance as the dependent variables.

## 2 Materials and methods

### 2.1 Cattle manure collection

The study was conducted on 50 Polish Holstein-Friesian dairy cows, Black and White variety. The herd was located in the central part of Poland (Masovian voivodeship). The animals were kept in a loose barn with free access to water, and fed the same total mixed ration (TMR) diet ad libitum, consisting of corn silage (75%), concentrates (20%), and hay (5%), supplemented with a mineral and vitamin mixture, according to the INRA system (ruminant feeding system developed at the Instytut National de la Recherche Agronomigue, France) (Brzóska et al., 2014). The cattle manure samples were collected in October 2019. The manure samples were deposited into two plastic drums measuring approximately 20L each. One aliquot of manure was subjected to composting, and the other to storage on a laboratory scale. The farm owner agreed to the manure being sampled and shared the history of the use of antibiotics on the farm.

### 2.2 Manure management strategies

Defined amounts of cattle manure (approx. 5 kg) were mixed with Sitka spruce (4:1 ratio; w/w) and placed in a bucket. The Sitka Spruce was sourced from a local hardware store. Containers with manure were placed in the laboratory under a ventilated hood at room temperature. All samples originate from the same batch of animal feces, collected during one sampling separated into two portions of manure - one designated for composting and second for storage. Each of them was in turn divided into three parts and each of them was mixed well with Sitka spruce (three biological replicates per treatment). Then each biological replicate was separated into three technical replicates. During composting, the manure was mixed well each week for ten weeks, while the stored manure was left undisturbed for four months. Although the duration of both processes differs, the composting and storage are both, highly recommended for manure treatment before its land application and both were applied according to commonly used practices - the composting process was completed within eight weeks (56 days), as noted by Mc Carthy et al. (2011) and the four-month storage period was selected based on EU legislation. Moreover, the same time intervals were applied by Do et al. (2022, 2023) during their study on manure treatment. Samples for the microbiological and molecular analysis were collected at the beginning (raw/untreated manure), in the middle (a fifth week for composting - 5W; second month for storage - 2M), and at the end of the process (a tenth week for composting - 10W; fourth month for storage - 4M). For statistical analysis purposes, samples were divided into five groups representing processes and time points; corresponding technical replicates were collected and pooled together as replicates. A total of 200 samples were collected during the entire experiment. From each group, we gathered 40 samples, encompassing three technical repetitions (represented as pooled samples from technical replicates). The groups included control samples, composted samples at five and ten weeks, and stored samples at two and four months. DNA isolated from these samples was grouped accordingly and then dispatched for analysis.

The pH, moisture, and temperature were measured in both mixtures at approximately half of their height (∼ 15 cm) each week for ten weeks (enough time to observe the dynamics of changes during composting and to stabilize parameters during storage). The moisture content was adjusted to 50-60% for compost by adding sterile MilliQ water. The moisture, pH and temperature was measured directly in the containers with treated samples along with the time. The moisture and pH was measured by a dual-purpose pH meter dedicated to environmental samples (pH-meter Stelzner 3000/Soiltester). Temperature was measured by laboratory thermometer (Testo 905-T1).

### 2.3 DNA extraction

The total DNA was extracted using the commercially available FastDNA™ SPIN Kit for Feces (MP Biomedicals, California, USA) according to the manufacturer’s recommendations. The quality and quantity of the extracted DNA was analyzed with a Qubit 4.0 Fluorometer using the dsDNA high-sensitivity assay kit (Invitrogen, Thermo Fisher Scientific, Waltham, MA, USA) and a Colibri spectrophotometer (Titertek Berthold, Pforzheim, Germany). The DNA samples were isolated in triplicate and then pooled to obtain a single DNA sample for each time point.

### 2.4 Sequencing the variable V3-V4 regions of bacterial 16S rRNA

The bacterial community structure was determined using Illumina technology on the Miseq platform by sequencing the variable V3-V4 regions of bacterial 16S rRNA. The libraries were prepared with Nextera® XT index Kit v2 Set A (Illumina, San Diego, CA, USA). Once prepared, amplicons underwent sequencing, 2x300 bp paired-end sequences. PCR and sequencing was performed at the DNA Sequencing and Oligonucleotide Synthesis Facility (Institute of Biochemistry and Biophysics Polish Academy of Sciences). Raw sequences were processed and analyzed using Qiime2 software suite (Bolyen et al., 2019) with the DADA2 option for sequence quality control and the newest release of the SILVA (SILVA SSU database 138, accession date October 2021) ribosomal RNA sequence database for taxonomy assignment (Quast et al., 2013; Yilmaz et al., 2014).

### 2.5 High-throughput qPCR

High-throughput qPCR was performed by an outside firm (Resistomap, Helsinki, Finland) using the qPCR SmartChip Real-Time PCR cycler (Takara; Kusatsu, Japonia). The qPCR cycling conditions and initial data processing were carried out as previously described by Wang et al. (2014). The analysis used a 384-well template of primers. All samples used for the analysis met the following criteria: 260/280 ratio of 1.8-2.0 (+/-0.1) and a concentration of 10 ng/μl.

The threshold cycle (Ct) values were calculated using default parameters provided by the SmartChip analysis software. All samples for each primer set underwent qPCR efficiency and melting curve analysis. Amplicons with unspecific melting curves and multiple peak profiles were considered false positives and excluded from further analysis. Samples meeting the following criteria were selected for analysis: (1) Ct ≤ 27, (2) at least two replicates, (3) amplification efficiency 1.8 - 2.2. The relative copy number was determined using an equation originally published by Chen et al. (2016) and adapted to our study conditions. The gene copy numbers were calculated by normalizing the relative copy numbers per 16S rRNA gene copy numbers.

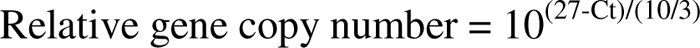

The genes and their corresponding primer sequences can be found in Supplementary Materials File 1. The gene names and groupings, as detailed and organized in the table from Supplementary Materials File 1, are consistently referred to throughout the manuscript.

### 2.6 Statistical analysis

The minimal sample number was determined by G*Power software (version 3.1.9.7) (Faul et al., 2007, 2009). The metagenomic data, including ARGs distribution, was analyzed and visualized using the MicrobiomeAnalyst web server (https://www.microbiomeanalyst.ca/)(Chong et al., 2020). Data were not rarefied, not scaled and transformed. The differences in microbial community structure were evaluated using the Qiime2 pipeline, based on the Kruskal-Wallis H-test (Shannon and Chao1 indexes). The differences between ARGs groups at each stage of the process were calculated using analysis of variance (ANOVA) with post hoc Tukey’s test (GraphPad Prism 9). The correlations between ARGs and MGEs and phylum and ARGs groups were calculated using Spearman’s correlation coefficient with GraphPad Prism 9. The networks representing correlations were built with Cytoscape (3.9.0 version) (Shannon et al., 2003). The correlations between ARGs and MGEs were considered strong and significant when the absolute value of Spearman’s rank |r|>0.9 and p<0.05 (Zhu et al., 2017), and for phylum and ARGs groups - |r|>0.9 and p<0.05.

## 3 Results

### 3.1 The changes in physicochemical properties of treated manure

The humidity, pH, and temperature were recorded weekly over a ten-week period, as detailed in Table S1 of Supplementary File 2.

### 3.2 Microbiome composition

During the analysis we identified 28 operational taxonomic units (OTU) at phylum level (Table S2 Supplementary Material File 2 - mean abundance with standard deviation of each phylum across the treatment). In all samples, the microbial community composition was determined for the untreated (raw) cattle manure and the composted and stored manure; these determinations were performed at the beginning, middle and end of the process. The composition of bacterial communities at the phylum level is presented in Fig. 1. using the taxon bar plot (additional table S1 in Supplementary material file 2), and the core microbiome is presented in Fig. S1 (Supplementary material file 2). In raw manure Bacillota has the highest abundance, while at the end of both processes the shift to Pseudomonadota as the most abundant phylum was observed. Interestingly, when physicochemical parameters of raw material changed along with both processes, new phyla were detected. During composting, Myxococcota, Bdellovibrionota, Chloroflexota, Gemmatimonadota, Acidobacteriota, Hydrogenedentes, Armatimonadota, WPS-2, Dependentiae, Abditibacteriota, and Deferribacterota increases its abundance, and become detected by applied method, while Euryarchaeota and Elusimicrobiota abundance decreases, reaching levels below detection limit. During storage, Myxococcota, Bdellovibrionota, Chloroflexota, Gemmatimonadota, Acidobacteriota become detectable, while Spirochaetota, Desulfobacterota, Elusimicrobiota prevalence becomes below the detection level. The richness and diversity of the microbial communities were determined using the Shannon (comparable results, p-value 0.08) and Chao1 (higher value for composted samples, p-value 0.025) indexes (Fig. S2, S3 Supplementary Material File 2). The raw data obtained from sequencing was uploaded to the Sequence Read Archive (SRA) repository (BioProject ID: PRJNA1007935) (NCBI). The PCoA of taxon abundance in all sample groups, calculated based on the Bray–Curtis distance, revealed significant visual separation between the initial (untreated) and treated samples (p-value<0.01) (Fig. S4 Supplementary Material File 2; the results of pairwise PERMANOVA analysis are presented in Table S2 Supplementary Material File 2).

**Figure 1.**
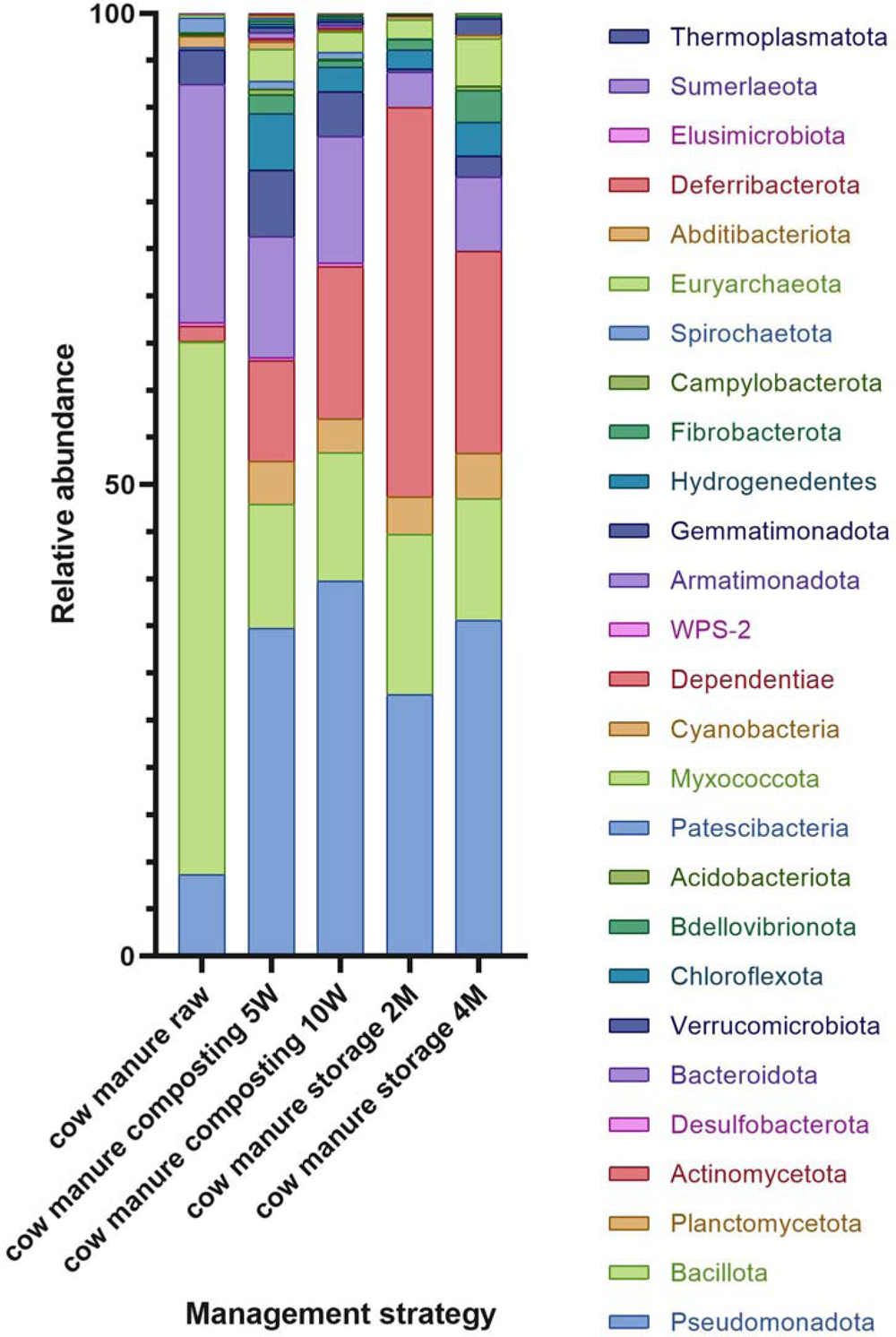
Relative abundance of phyla across samples; the samples were divided into five groups– control (cow manure raw; sample before treatment), cow manure composting 5W, cow manure composting 10W (composted samples after five weeks and ten weeks, respectively), cow manure storage 2M and cow manure storage 4M (stored samples after two months and four months, respectively); stacked barplots represents mean value for replicates.

### 3.3 Diversity and abundance of ARG

During the study, we found 202.67 (mean value from replicates) analyzed genes (Aminoglycoside-, Beta-Lactam-, cross-resistance to macrolides, lincosamides and streptogramin B (MLSB), Phenicol-, Sulfonamide-, Tetracycline-, Trimethoprim-, Vancomycin-resistance genes together with Integrons, MDR pumps (multi-drug resistance), MGEs, and ‘Other’ genes) in raw manure, 76 in composted samples after five weeks, 54 after ten weeks, and 51 in stored samples after two months and 39 after four months. The relative abundances of the analyzed ARGs classes and MGEs varied as storage and composting processed (Fig. S4 Supplementary Material File 2). Specifically, the relative prevalence of beta-lactam-, MLSB- and vancomycin-resistance genes fell in the treated manure samples compared to the untreated samples. In contrast, the relative abundance of genes encoding MGE, integrons and sulfonamide-resistance genes increases, but only after composting. However, the total gene count (i.e. total resistance gene abundance normalized per 16S rRNA gene) rapidly decreased following the application of both treatment strategies. Notably, for composting, the gene relative abundance was even lower in the middle of the process than at the end (Fig. 2).

**Figure 2.**
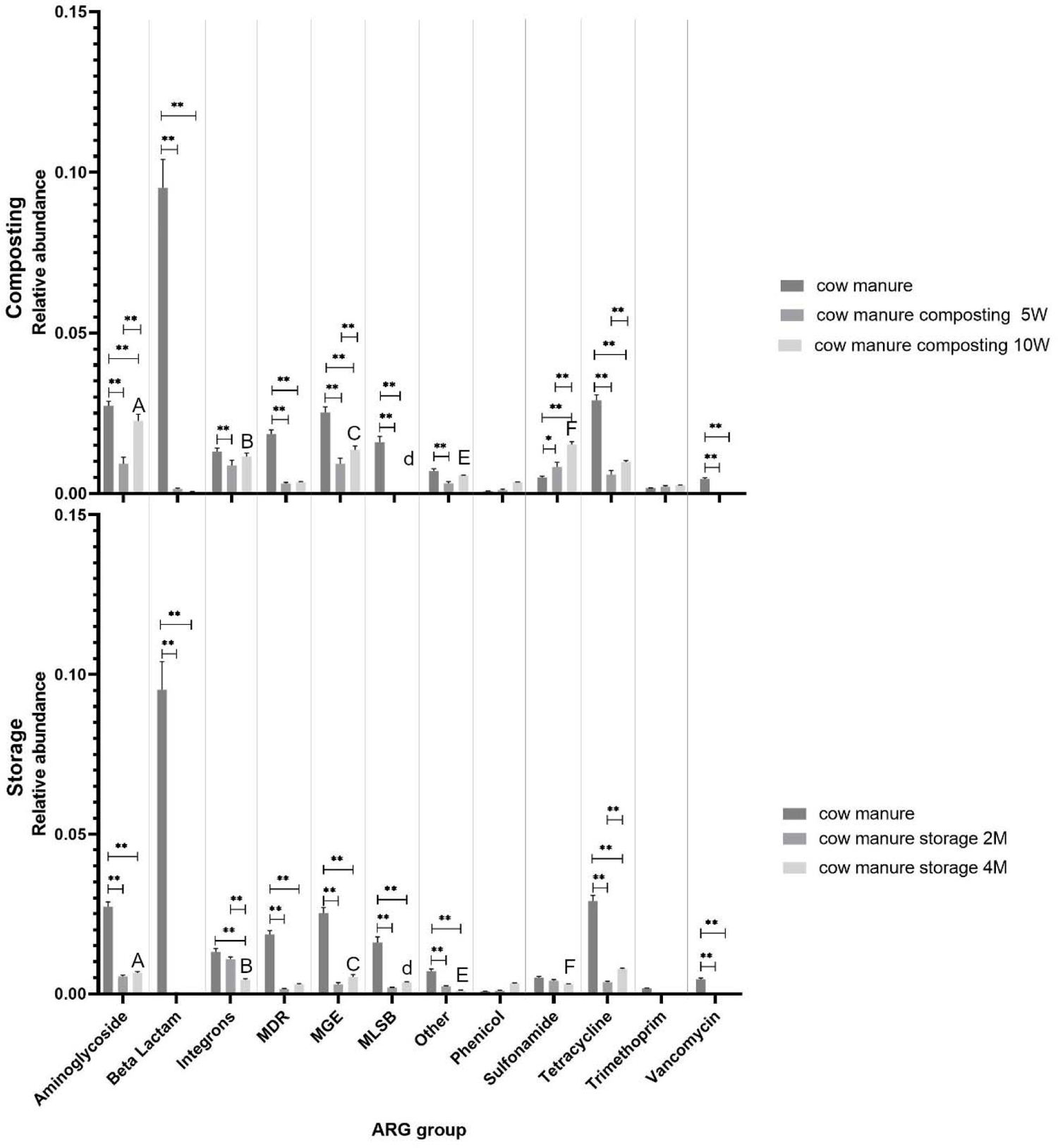
The differences in the amounts of ARGs and MGEs detected during composting and storage compared with raw manure. The following cut-off points for significance within the same process were applied: the values differ significantly at pL<L0.01 (indicated as *); the values differ significantly at pL<L0.05 (indicated as **); the values do not differ significantly at pL>L0.5. Relative abundances of genes were additionally compared at the end of both processes between them; the value with the same letters differs significantly: A, A – at p<0.01.; a, a at p<0.05 (indicated as A, B, C, D, E, F); the values differ significantly at pL<L0.05 (indicated as d).

Composting efficiently reduced the quantity of aminoglycoside- and tetracycline-resistance genes, as well as MGEs, within the initial five weeks (2-fold, 5-fold, 2.5-fold reduction respectively, p<0.01). However, a subsequent rise in their abundance was observed after ten weeks of treatment compared to their level after five weeks of treatment. Overall, compared to raw material it still remains lower (1,2-fold, 3-fold and 2-fold reduction respectively, p<0.01). Beta-lactam-, vancomycin-resistance genes, MLSB-coding genes, and genes responsible for MDR demonstrated a decline at both the five-(66-fold for beta-lactam-resistance genes, 74 for MLSB-coding genes, and 6-fold for MDR genes, with vancomycin-resistance genes being below detection limit; p<0.01) and ten-week (156-fold for beta-lactam-resistance genes, 80-fold for MLSB-coding genes, and 5-fold for MDR genes, with vancomycin-resistance genes being below detection limit; p<0.01) marks, with the rate of reduction being consistent between the two points. The abundance of Integron-coding genes decreased after five weeks of composting (1.5-fold, p<0.01), but remained unchanged after ten weeks compared to raw sample and sample after five weeks of composting. Neither phenicol-nor trimethoprim-resistance gene abundance varied between composting stages. However, the sulfonamide-resistance gene count increased after five weeks of composting (0.6 fold, p<0.05), and exhibited an even greater increase after ten weeks (0.33-fold, p<0.01).

Our results show that storage effectively diminishes the quantities of aminoglycoside-resistance genes (5-fold, p<0.01), MGEs (9-fold, p<0.01), and genes responsible for MDR (12-fold, p<0.01) and MLSB-coding genes (8.5-fold, p<0.01) after two months. However, their numbers subsequently increased after four months of treatment, but still remains lower than in raw manure (p<0.01). The counts of Beta-lactam, vancomycin-resistance genes, and those labeled as ‘Other’ demonstrated a decline both at the two-(289-fold, 8.5-fold, respectively, with vancomycin-resistance genes being below detection limit; p<0.01) and four-month marks (579-fold, 7-fold, respectively, with vancomycin-resistance genes being below detection limit; p<0.01), yet the rate of reduction remained consistent between these points. The abundance of genes coding for integrons remained consistent after two months of storage but declined after four months (3-fold, p<0.01). No noticeable change in the amounts of phenicol-, sulfonamide-, and trimethoprim-resistance genes was observed across the storage periods. The tetracycline-resistance genes were lower after both the two (8-fold, p<0.01) and four month (4-fold, p<0.01) periods, but their concentration was notably higher at the end of the fourth month compared to the second.

Both composting and storage processes reduce the abundance of aminoglycoside-resistance genes, with storage achieving a greater reduction. A similar trend can be observed for the abundance of MLSB-coding genes, MGEs, and genes classified as ‘Other’. For beta-lactam, tetracycline-, vancomycin-resistance genes, and genes responsible for MDR, a reduction in gene abundance was noted after both processes. However, composting and storage yielded comparable outcomes by the end of treatment. While storage resulted in a decreased abundance of integron-coding genes relative to untreated samples and ten-week composted manure, composting itself did not appear to affect these genes. For sulfonamide-resistance genes, composting increased their amount, while storage did not include any noticeable changes. Nevertheless, after two months of composting, the abundance of sulfonamide-resistance genes remained elevated compared to post-storage levels. The quantities of the phenicol- and trimethoprim-resistance genes did not demonstrate any significant variation after the two treatments compared to untreated manure (Fig. 2).

The relative abundances of ARGs and MGEs across all analyzed samples were subjected to principal coordinate analysis (PCoA), using the Bray-Curtis distance method. The results indicate a significant visual separation between the untreated manure (control) and treated samples with regard to the relative abundances of resistance gene classes (p<0.01).

The core resistome across all analyzed samples consists of 29 genes. They belong to the following groups: aminoglycoside-resistance genes (*aadA5_1, aadA_2, aac3-VI, aacC2, aadA1, strB, aadA2_3, aadA2_1*), beta-lactam-resistance genes (*fox5*), phenicol-resistance genes (*floR_1*), sulfonamide-resistance genes (*sul1_1, sul2_1*), tetracycline-resistance genes (*tetA_2, tetG_1, tetG_2*), trimethoprim-resistance genes (*dfrA1_1, dfrA1_2*), integron-coding genes (*intI1_3, intI1_1, intI1_4*), genes responsible for MDR (*mdtH_1, oprJ, acrR_3*), MGE (*tnpA_2, ISSm2, IncP_oriT, repA*), and resistance genes classified as ‘Other’ (qacEΔ1_2, qacEΔ1_3). For both applied treatment strategies, a decreased number of ARGs were observed post-treatment compared to raw manure (53 for compost and 38 for storage) (Fig. S5 Supplementary material file 2).

In all composted and stored samples, almost all ARGs, excluding trimethoprim-resistance genes, were found to be reduced at the end of both processes, which means lower diversity of genes after treatment. In addition, the stored samples demonstrated lower numbers of aminoglycoside-, beta-lactam-, phenicol-, sulfonamide-, tetracycline-resistance genes and MGEs compared with composting. However, the composted samples demonstrated lower numbers of genes conferring MLSB and MDR resistance compared to the stored samples (Fig. 3).

**Figure 3.**
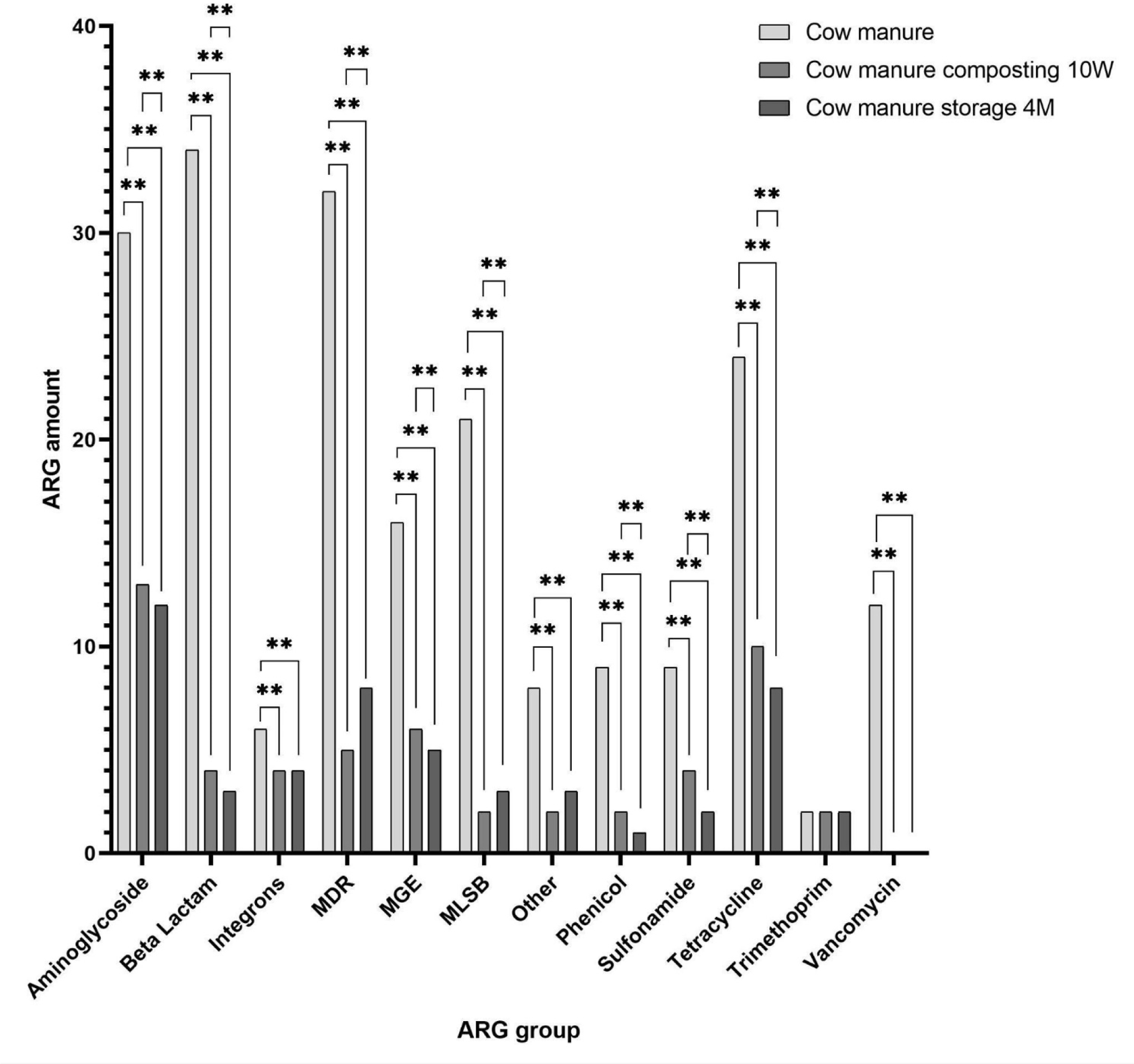
Comparison between ARGs (assigned to ARGs classes) at the end of both treatment processes.

In the composted samples, the correlation network analysis revealed 124 nodes connected by 512 edges, representing strong and significant correlations (|r|>0,99; p<0.05) (Fig. 4). All correlations were positive. The MGEs (integrons included) exhibited more links than ARGs, indicating their significant role in network formulation. The MGEs (n=15) included in this network are *intI3_1, orf39_IS26, IncN_rep, Tn5, Tp614, tnpA_5, ISAba3, trfA, IS613, intI3_2, tnpA_6, intI1_3, ISSm2, IS1111, tnpA_2*.

In the stored samples, strong and significant correlations were found between MGEs (integrons included) and ARGs levels, manifested as a network containing 144 nodes connected by 643 edges (|r|>0.99; p<0.05) (Fig. 5). Every correlation in the network was positive. The MGEs (integrons included) had more connections than ARGs, indicating they play a pivotal role in the formation of the network. The MGEs (n=16) presented in the network are *intI3_2, IS613, IS1111, ISAba3, orf37-IS26, tnpA_1, trfA, tnpA_5, orf39_IS26, Tn5, Tp614, tnpA_6, intI2_2, IncN_rep, intI1_1, intI3_1*.

**Figure 4.**
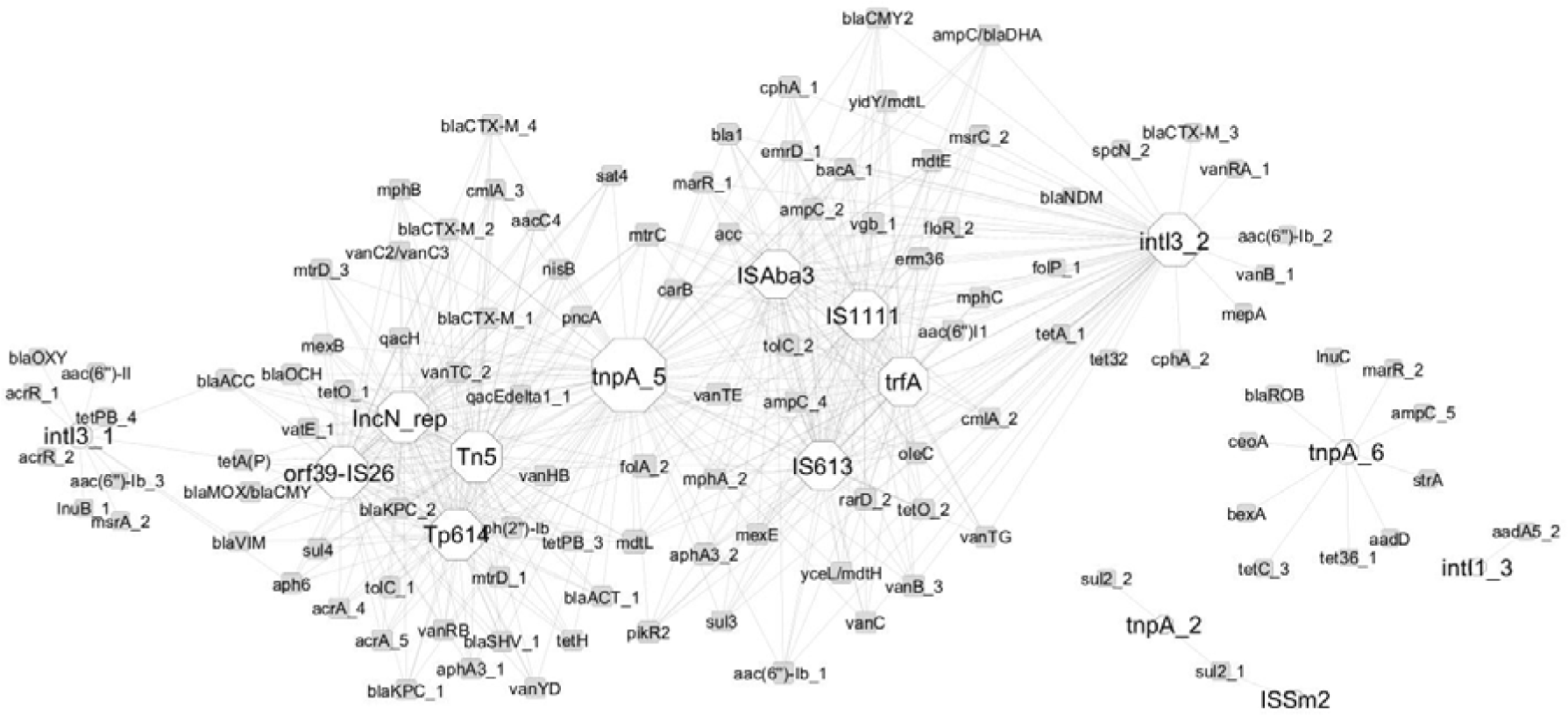
ARGs and MGEs interaction network in composted manure. The network is presented as an ‘organic layout’. Based on Spearman’s rank correlation, a strong and significant correlation is shown, where |r|>0.99 and p < 0.05. The size of the nodes represents the degree of interaction. The gray edges show positive correlations between ARGs and MGEs. The color of the nodes indicates ARG (white) and MGE (gray).

**Figure 5.**
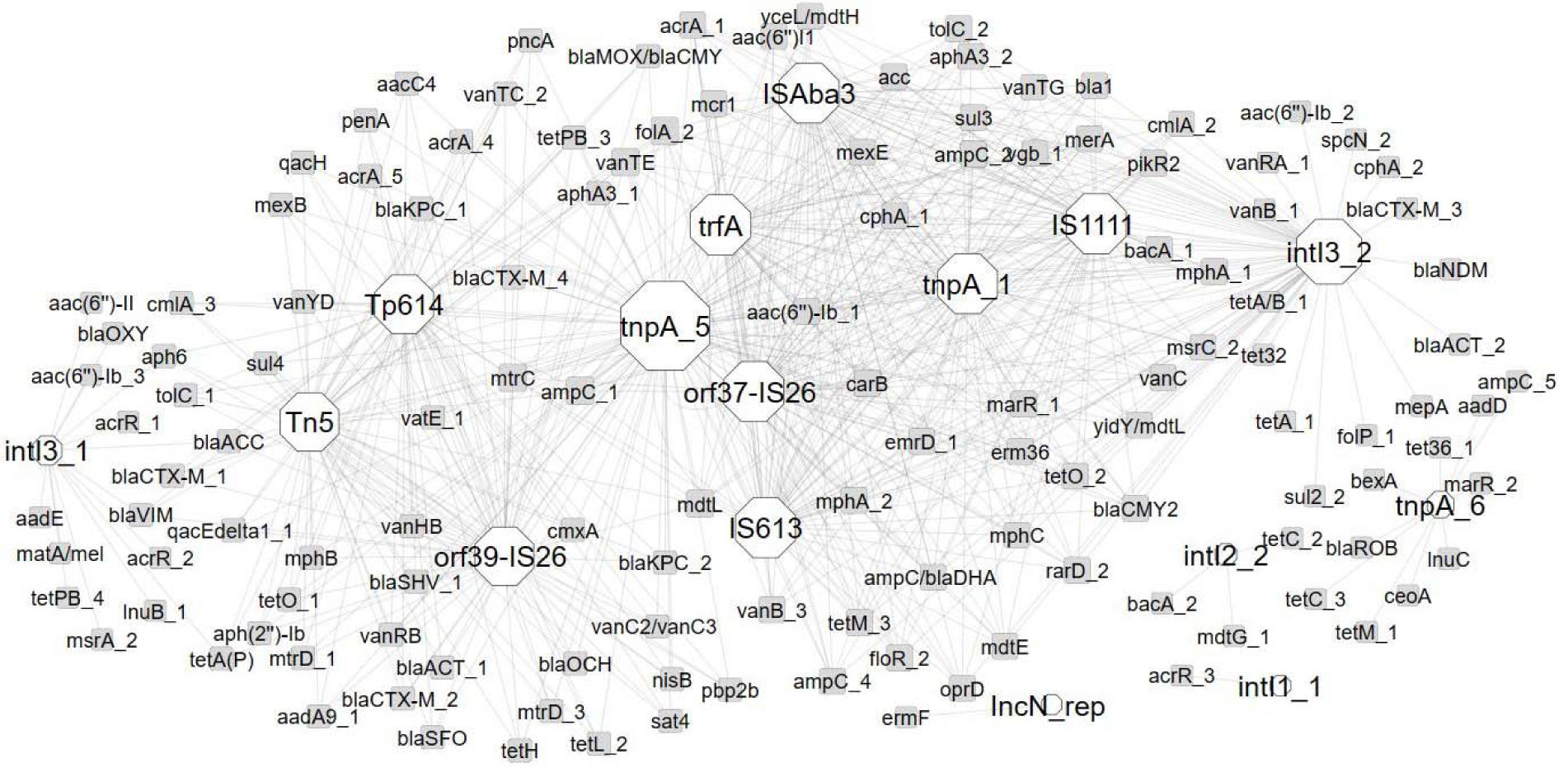
ARGs and MGEs interaction network in stored manure. The network is presented as an ‘organic layout’. Based on Spearman’s rank correlation, a strong and significant correlation is shown, where |r|>0.99 and p < 0.05. The size of the nodes represents the degree of interaction. The gray edges show positive correlations between ARGs and MGEs. The color of the nodes indicates ARG (white) and MGE (gray).

In composted samples, negative correlations were found between beta lactam resistance genes and MDR and Spirochartota and Cyanobacteria, and positive ones between MLSB and Acidobacteriota and WPS-2 group (Fig. 6).

**Figure 6.**
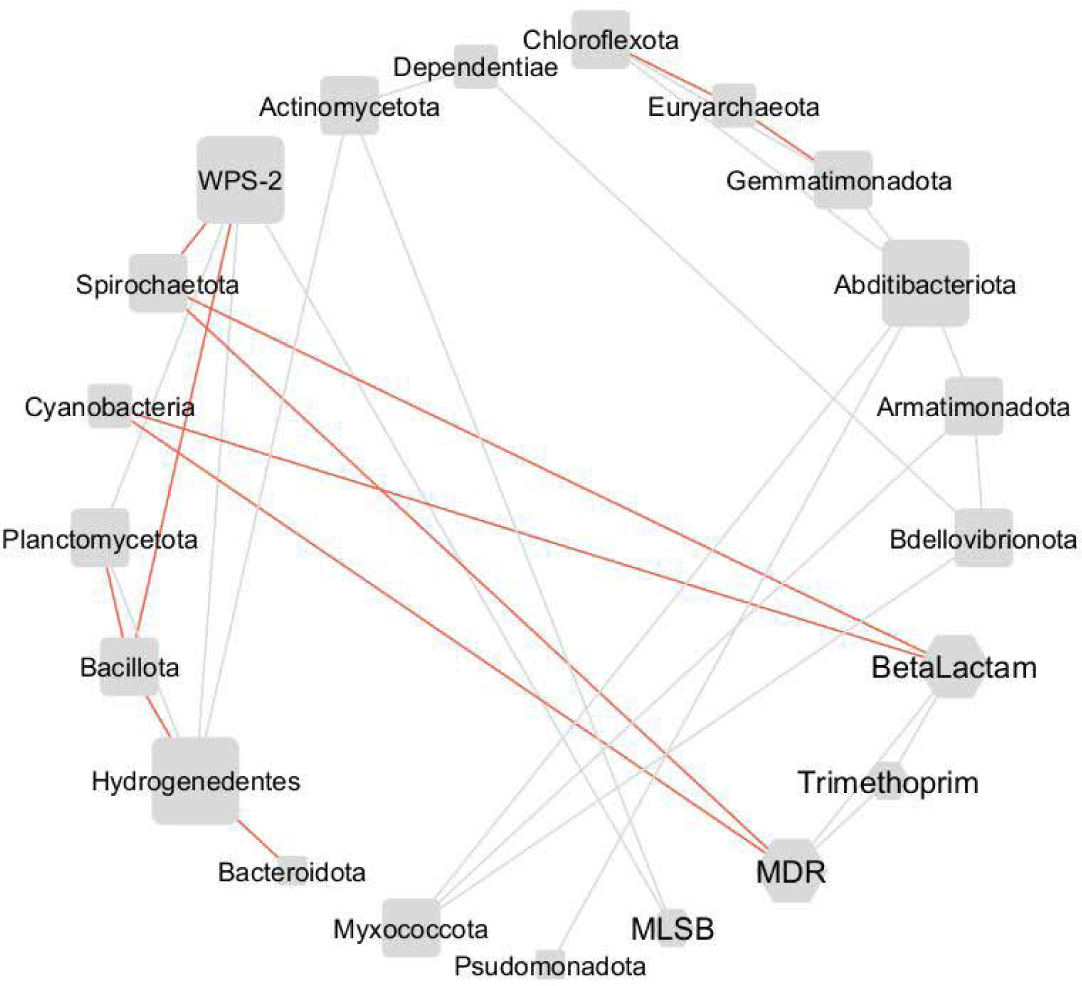
Phylum and ARGs group interaction network in composted manure. The network is presented as a ‘circular layout’. Based on Spearman’s rank correlation, a strong and significant correlation is shown, where |r|>0.9 and p<0.05. The size of the nodes represents the degree of interaction. The gray edges show positive correlations, while red one - negative. The shape of the nodes indicates phylum (square with rounded corners) and ARGs group (hexagon).

In stored samples, only negative correlations were identified: between sulfonamide and Bdellovibrionota, Chloroflexota and Gemmatimonadota, between tetracycline and Bdellovibrionota, Gemmatimonadota, Chloroflexota and Myxcoccota, between vancomycin and Desulfobacterota and between aminoglycosides and Campilobacterota; however, vancomycin and aminoglycosides creates separate clusters (Fig. 7).

**Figure 7.**
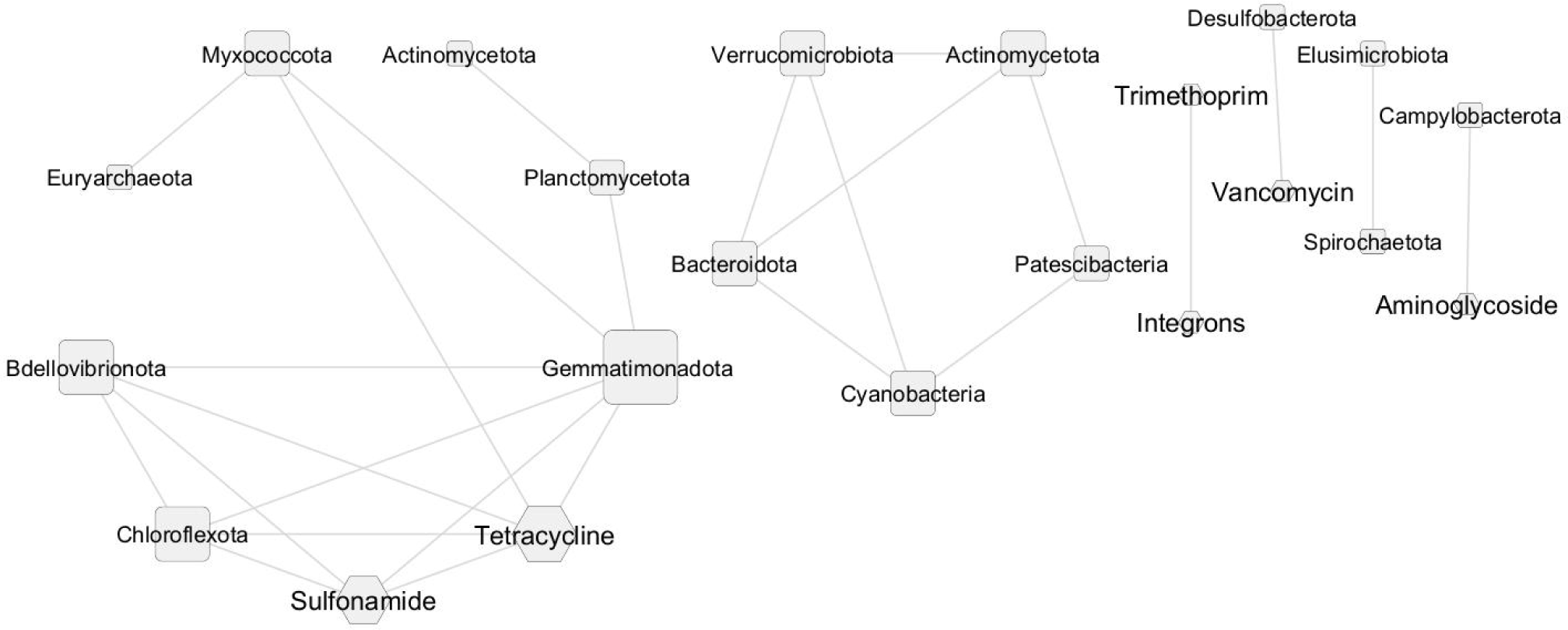
Phylum and ARGs group interaction network in stored manure. The network is presented as a ‘circular layout’. Based on Spearman’s rank correlation, a strong and significant correlation is shown, where |r|>0.9 and p<0.05. The size of the nodes represents the degree of interaction. The gray edges show positive correlations, while red one - negative. The shape of the nodes indicates phylum (square with rounded corners) and ARGs group (hexagon).

## 4 Discussion

Farms often dispose of animal manure by direct application to the land. Many also employ a range of manure management technologies to reduce environmental pollution before application. These technologies not only reduce the total manure volume, but also can be used to produce organic fertilizer, or possibly facilitate biogas production. Two such technologies are storage and composting, after which the feces are often used as organic soil amendments with potential benefits for soil quality and crop yield (Epelde et al., 2018). Such amendments typically contain all nutrients essential for plant growth and can replace or complement inorganic fertilizers in agricultural production systems (Youngquist et al., 2016). However, cattle manure can also serve as a reservoir of environmental ARG pollution (Udikovic-Kolic et al., 2014; Zalewska et al., 2021), and the direct use of livestock manure without proper treatment could promote the dispersal of ARG in arable soil and cultivated plants (Ghosh and LaPara, 2007; Heuer et al., 2011; Udikovic-Kolic et al., 2014; Zalewska et al., 2023). Manure treatment strategies have not been specifically designed to mitigate AMR (Oliver et al., 2020).

Our present findings indicate that composting and storage are effective methods of reducing ARGs from manure before use. It is hard to define which process accomplishes this task better, as despite a generally-observed significant reduction in the number of ARGs, particular groups’ of resistance genes responded differently to the applied strategies. For example, the levels of aminoglycoside-resistance genes, integrons and MGEs, genes classified as ‘other’ and sulfonamide-resistance genes were lower after storage than composting. In contrast the MLSB levels in the stored samples were higher than those in the composted samples, but they were still lower than those found in the raw material. No differences were observed between the composted and stored samples for beta-lactam, MDR, tetracycline, and vancomycin-resistance genes; those genes are often prevalent in animal manures. Similar results for composting were obtained by Gou et al. (2018), who found higher amounts and numbers of ARGs in unprocessed manure compared to composted samples.

### 4.1 Bacterial community structure

Wang and Zeng (2018) reported that bacteria play a greater role in composting than fungi. Pitta et al. (2016) identified several prevalent bacterial phyla in dairy cattle feces and raw manure: Bacterioidota, Bacillota, Pseudomonadota, Actinomycetota, Acidobacteriota and Spirochaetota. This aligns with our present findings, which also included Spirochaetota and Acidobacteriota, though they were not the most represented phyla. Moreover, Zhou and Yao (2020) report the presence of Pseudomonadota and Bacillota as the main phyla in animal feces.

The decomposition of organic matter starts before composting commences, as microorganisms are introduced unintentionally to the organic matter during grinding, mixing, cutting, and transporting. The composting pile initially rests at an ambient temperature. It then gradually heats up during the mesophilic phase to 25-45 °C, indicating increased microbial activity (Ezugworie et al., 2021). The diversity of microbial community correlates with the metabolic functions of the microbes, e.g., lactic acid-producing bacteria and acetate-oxidizing bacteria are usually detected at the mesophilic phase of the composting process, where they utilize the easily degradable and soluble organic matter. Zhang et al. (2022) have found that Bacillota, Pseudomonadota, Bacteroidota, Chloroflexota, Actinomycetota, and Planctomycecota were the dominant phyla during co-composting of cow manure and maize straw. In addition, Zhang et al. (2020) report the presence of Bacillota, Bacteroidota, Deinococcota, Pseudomonadota, and Actinomycetota in raw manure and composted samples; however, the abundance of Deinococcota and Actinomycetota were much higher in composted samples. Similarly, Zhou and Yao (2020) noted that composting resulted in a shift in the bacterial community, with an increased prevalence of Actinomycetota compared to raw manure. Our present findings also identified Pseudomonadota, Bacillota, Planctomycetota, and Actinomycetota as being the most prevalent during composting; however, their ratio changed with time. Zhang et al. (2016) found the Bacillota, Pseudomonadota, Bacteroidota, and Actinomycetota to predominate in the thermophilic phase of composting. Of these, Bacillota were the dominant phylum due to their adaptability to aerobic condition and heat resistance. Temperatures exceeding 70 °C terminate most microbial activities, forcing certain bacteria to sporulate (Ezugworie et al., 2021).

While our sequencing results differed from those found in some previous studies, the identified phyla remained consistent. Composting can be influenced by a range of physicochemical factors, such as temperature, moisture, pH, aeration, and carbon-to-nitrogen ratio. As such, as there are no strict guidelines on composting, composted samples can differ widely. A crucial role is played by temperature: it governs the pace of biological processes conducted by microorganisms and determines the presence of particular microorganisms; for example, fungi, which are not heat-resistant, are eliminated at temperatures above 50 °C. In addition, a pH of 7-8 is needed for optimal composting, which slows down at low pH. The C/N ratio should also be kept at optimum, as nitrogen is crucial for microbial growth, and carbon is an energy source. Humidity should be kept at 40-60% to ensure the transfer of dissolved nutrients necessary for microbial growth (Ezugworie et al., 2021). As such, the differences in bacterial community structures obtained by different research teams may hence be due to the differences in physicochemical parameters during the process. In our case, the temperature was within the proper range, but the humidity and pH were slightly suboptimal (humidity above 80% and pH below 7).

### 4.2 Changes in antibiotic resistance genes after treatment

#### 4.2.1 Tetracycline-resistance genes

The tetracyclines are the most commonly used antibiotics in the livestock production sector; as such, tetracycline-resistance genes are the subject of intensive study. These genes appear to be unaffected by manure treatment strategies such as composting or storage, and the detection rate of t*etW* and *tetO* may account for up to 80% of fecal samples obtained from cattle raised in grassland-production systems, even without antibiotic usage (Santamaría et al., 2011). The following tetracycline-resistance genes were identified in the present study: efflux (*tetA, tetA/B, tetG, tetH, tetC, tetL, tetA(P)*), protection (*tet(36), tet(32), tetO, tetQ, tetW, tetM, tetPB*), deactivation (*tetX*) and regulation (*tetR*) in raw manure samples. Our findings are partially in line with those of other studies (Jauregi et al., 2021). Zhang et al. (2020) found *tetW, tet40, tetO, tetQ*, and *tetC* to be the most prevalent types across all analyzed raw dairy cattle manure samples, accounting for over 40% of detected ARGs. Marti et al. (2013) report the presence of *tetM, tetQ, tetS, tetT, tetA, tetB/P* and *tetW* in cattle manure, and Mu et al. (Mu et al., 2015) found *tetM, tetO, and tetW* in cattle manure, and *tetM, tetO, tetQ*, and *tetW* in feces collected per rectum; the latter partially agrees with our present findings indicating these genes to be present in manure.

However, Zhou and Yao (Zhou and Yao, 2020) reported a general increase in the abundances of total tetracycline-resistance genes, which contradicts our results. Despite this, as in our present study, Zhang et al. (Zhang et al., 2020) failed to detect *tetA, tetW*, and *tetR*, and that total ARGs abundance decreased after composting. Interestingly, Zhang et al. (Zhang et al., 2020) also indicate a reduction in *tetW, tetA*, and *tetR* abundance with composting time, together with a decrease in the concentration of the total amount of ARGs. Staley et al. (Staley et al., 2021) did not identify any differences in the concentrations of *tetO* and *tetQ* genes between composted and stored samples; however, their level was found to change with sampling season and depth. In addition, Qian et al. (Qian et al., 2016) report increased abundance of *tetC* and *tetX*, and decreased abundance of *tetQ, tetM* and *tetW* after composting.

Interestingly, while Wang et al. (Wang et al., 2019) did not identify any decrease in *tetW, tetO* or *tetM*, Storteboom et al. (Storteboom et al., 2007) indicate that the *tetO* and *tetW* genes appeared to gradually decrease during composting; this agrees with our present findings indicating that *tetO* was present in the middle of composting, but below the detection limit at the end of the process. However, our data indicate that *tetW*, together with *tetA, tetG, tetPB, tetQ, tetM, tetR, tetX,* and t*etA/B*, remained persistent during composting.

Another study examining the tetracycline-resistance genes *tetO* and *tetA* found no noticeable change in their abundance, even after one year of storage (Hurst et al., 2019). These findings are in line with ours, where *tetA* remained persistent throughout storage, contrary to *tetO*. Interestingly, Jauregi et al. (Jauregi et al., 2021) found *tetW, tetO, tetM, tetB, tet32, tetN*, and *tetG* to be present in stored cattle manure even after six months; in contrast, *tet32, tetO, tetW,* and *tetM* were below the detection limit after two months of storage but reappeared after four months.

Tetracycline-resistance genes can be transmitted to gram-negative and gram-positive bacteria species, e.g., *tetM, tetW*, and *tetQ* originate from gram-positive bacteria and can be transmitted across Gram-positives and Gram-negatives (Chopra and Roberts, 2001). Moreover, tet genes are often placed on plasmids or inserted in transposons, facilitating their dissemination (Lima et al., 2020). Jauregi et al. (Jauregi et al., 2021) regard *tetM, tetO, tetW, tet32*, and *tetS* as being of particular concern due to their high dissemination potential associated with their co-occurrence with MGEs; they also highlight *tet44* and *tetW/N/W*, which were not detected during our study. Our results partially agree, as our data also indicates some tetracycline-resistance genes to be significantly associated with MGEs: *tetM (tnpA, IS613, trfA, orf37_IS26, ISAba3, IS1111, IntI3), tetO (tnpA, trfA, orf39_IS26, orf37_IS26, ISAba3, IS1111, Tn5, IntI3, tp614, IS 613), tet32 (intI3)*; however, neither *tetW* or *tetS* were confirmed in samples obtained from either process. We also found *tet36, tetPB, tetC, tetA(P), tetC, tetH,* and *tetL* to have a high probability of being exchanged by horizontal gene transfer (HGT) due to co-occurrence with MGEs.

#### 4.2.2 Beta-lactam-resistance genes

The following beta-lactam-resistance genes were identified in raw manure, classified as deactivation (*ampC, blaMOX/blaCMY, blaOCH, blaSHV, blaROB, blaOXY, cphA, cfxA, blaCMY2, ampC/blaDHA, blaFOX, blaCTX-M, blaSFO, blaVIM, blaKPC, blaOXA, blaNDM, blaACC, bla1, blaCMY, blaACT*) and protection (*penA, pbp2b*). Our results are similar to those of Marti et al. (2013), who identified *blaOXA, blaCTX-M, blaPSE* and *blaVIM*, as well as *blaTEM*, which was not found in the present study. Pitta et al. (2016) report the presence of *pbp1A, pbp2*, and *pbp2B* in dairy cattle manure; however, these were not present in our samples. While Zhou and Yao (2020) confirm our observed general decrease in beta-lactam-resistance genes, Keenum et al. (2021) found the levels of beta-lactam-resistance genes to be elevated after composting.

Hurst et al. (2019) did not identify *blaOXA, blaCTX-M*, and *blaVEB-1* after long-term storage of dairy cattle manure. This is in line with our present findings, which did not identify any beta-lactam-resistance genes present in raw manure in the samples subjected to long-term storage and composting, except for the *fox5* gene; however, *blaVEB* was not observed in our raw manure.

The *blaTEM* and *blaCTX-M* genes are often associated with *IncN* plasmids, which are believed to play an essential role in beta-lactamase gene dissemination (Lima et al., 2020). Our study found *bla* genes to be significantly associated with *IncN_rep* in composted manure samples but not in stored ones. Moreover, beta-lactam-resistance genes are often located in insertion sequences such as *ISEcp1* or *ISCR1* (Lima et al., 2020). Although our findings did not identify the insertion sequences mentioned above, they did indicate the presence of other MGEs associated with *bla* genes viz. *IS613, trfA, ISAba3, IS1111, intI3, tnpA, Tp614, orf39-IS26, Tn5, IS613*; this suggests they have high mobility in both treatment strategies.

#### 4.2.3 Aminoglycoside-resistance genes

The *aadA, aadA5, aadD, aadA9, aphA1/7, aac(6’)I1, aac3-VI, aacC2, aadA1, aadE, strA, strB, acc, aacC4, aph6, aac(6’)-Ib, aadA2, aph(2’)-Ib, aphA3, aadA9, spcN, aac(6’)-II, aphA3* genes were observed in the present study, all of which are responsible for deactivation. Zhou and Yao (2020) report a general decrease in aminoglycoside-resistance genes during treatment, which agreed with our results.

Hurst et al. (2019) also found *aac-(6)-lb* and *aada1* to be present in manure after long-term storage; however, our present findings indicate that *aadA1* was also persistent in manure after storage as well as composting, but *aac(6)-lb* was not detected after either process. Although *aadA1* persisted after composting, its concentration was reduced, as also noted by Zhang et al. (2020).

Moreover, Jauregi et al. (2021) found *aad6* and *ant6-lb* to be of particular concern due to their co-occurrence with MGE and hence their high dissemination potential. However, although these genes were not identified in raw manure in the present study, other genes of potential concern (i.e. co-occurring with MGE) were noted: *aac(6’’)-Ib (intI3, IS613, tnpA, trfA, orf37-IS26, ISAba3, IS1111, intI3), aac(6’’)-Ib (intI3, IS613, tnpA, trfA, orf37-IS26, ISAba3, IS1111) and aphA3 (Tp614, IS613, tnpA, trfA, orf37-IS26, orf39-IS26, ISAba3, IS1111, Tn5, intI3*).

#### 4.2.4 MLSB-resistance genes

Despite their wide application in livestock, often with lincosamides and streptogramin, the data on resistance genes against macrolides is limited. Mu et al. (2015) identified *ermB* and *ermC* in cattle feces (collected per rectum) and manure; however, the study only addressed macrolide-resistance genes. Our present data did not confirm the presence of *ermB* and *ermC*, but did identify other genes coding MLSB resistance phenotypes in cattle manure, such as *ermK, ermF, ereB, lnuB, vatE, erm36, matA/mel, mphA, mphB, vgb, mefA, msrC, pncA, mphC, msrA, lnuC, oleC, carB*, and *pikR2*. In contrast, Wang et al. (2019) found *ermB* and *ermF* to be present in raw cattle manure, while Pitta et al. (2016) found *carA, mefA, tlrC*, and *vgaA* genes to also be present in feces and manure.

Zhou and Yao (2020) report a general decrease in MLSB-resistance genes in total, which agreed with our results. However, Hurst et al. (2019) found *ermB, ereC*, and *mefA* to remain persistent in manure after long-term storage; our present findings did not indicate the presence of *ermB* in dairy cattle manure, and *ereB* and *mefA* genes to be removed by long-term storage and composting. Wang et al. (2019) did not identify any changes in *ermB* and *ermF* gene abundances during composting, since their levels were too low at the beginning of the process.

Lima et al. (2020) report *ermB* to be frequently incorporated in the *Tn916-Tn1545* family of conjugative transposons. Although *ermB* was not found in the present study, other MLSB phenotypes were noted including genes co-occurring with MGE: *IS613, trfA, ISAba3, IS1111, intI3* and *tnpA* in compost, and *IS613, tnpA, trfA, orf37-IS26, ISAba3, IS1111, intI3* and *IncN_rep* in storage. The presence of *IncN_rep* in stored samples should be especially concerning because it suggests the location of erm genes on a broad-range host plasmid from the IncN group.

#### 4.2.5 Resistance genes against other antimicrobials

Although the tetracycline- and beta-lactam-resistance genes are considered to be most prevalent in dairy and beef cattle feces, Marti et al. (2013) found chloramphenicol-resistance genes to be predominant, with tetracycline-resistance genes in fourth place. The *cmlA* and floR genes had high dissemination potential, due to co-occurrence (Jauregi et al., 2021), but our data indicates that *cmlA* was efficiently eliminated from animal manure after storage or composting. However, we agree that the *floR* gene should be considered highly mobile as our findings indicate that it could be associated with MGEs such as *IS613, tnpA_1, trfA, orf37-IS26, ISAba3, IS1111* and *intI3*, in both analyzed treatment strategies.

The effect of composting on sulfonamide-resistance genes has also been analyzed. Zhou and Yao (2020) report a general decrease in sulfonamide-resistance genes, which was in line with our present results; in addition, Mu et al. (2015) also identified *sul1, sul2*, and *sul3* in cattle manure, as noted in the present study. Additionally, our study also identified *sul4, folP*, and *folA*. The presence of *sul1* and *sul2* was also confirmed (Wang et al., 2019); however, their study did not show a reduction in the abundance of those genes after composting.

Furthermore, being often located on transposable elements of self-transposable or mobilizable broad host-range plasmids, *sul* genes are highly prevalent in a wide range of bacterial species (Lima et al., 2020). The *sul2* gene should be especially concerning, because it is often embedded on small mobile plasmid of the IncQ family with a broad host spectrum (Wang et al., 2019). The *sul3* gene has also been found in many sources together with class 1 integrons (Lima et al., 2020). Our present findings indicate the *sul* gene family to be associated with *intI3, tnpA, Tp614, orf39-IS26, Tn5, IncN_rep, IS613, trfA, ISAba3, IS1111* in both processes; however, *intI1* was not noted during our study

In addition, *qnrA* and *qnrB* resistance genes were not identified in the present cattle manure samples. Similarly, Mu et al. (2015) were also not able to detect the sought quinolone-resistance genes (*qnrD* and *qnrS*). However, Wang et al. (2019) detected the presence of *qnrB* and *qnrS* genes in raw cattle manure, but did not observe any reduction after composting. Pitta et al. (2016) confirm the presence of *vanRA, vanRE*, and *vanRG* resistance genes in raw manure. While *vanRA* was found together with *vanB, vanC, vanC2/vanC3, vanHB, vanRB, vanTC, vanTE, vanTG, vanYB* and *vanYD* in the present study, *vanRE* and *vanRG* were absent. Finally, Zhou and Yao (2020) did not find any change in total vancomycin-resistance gene levels after composting, which contradicts our present results indicating a decrease in abundance.

Although significant reductions in ARGs abundances grouped according to antibiotic type (Fig. 2) were noted after composting and storage, the effect of those treatment strategies on particular genes is more complex, and many gaps still need to be addressed. The changes in ARGs abundance during storage or composting may vary depending on the location of the long-term storage pit or composting pile, storage/composting time (age), manure removal techniques (scrap versus flush removal) or even season of storage (Hurst et al., 2019). In addition, during composting, ARGs and MGEs removal can also be influenced by temperature, pH, moisture, or type of waste and bulking material (Pezzolla et al., 2021). In addition, ARGs levels can also depend on the co-occurrence of ARGs or MGEs, and on various factors affecting bacterial fitness, gene regulation and transfer, such as nutrient accessibility or heavy metal concentration (Oliver et al., 2020). While ARGs concentration also seems to be dependent on on-site pressure, such as the presence of antibiotics or heavy metals, it also appears to be influenced by the specific gene. Zhang et al. (2018) found lower levels of macrolide-resistance genes (*ermX, ermQ, ermF, ermT, and ermB*) to be associated with lower tylosin concentrations, due to the effect of tylosin degradation during composting. Guo et al. (2018) reported that adding cyromazine (a non-antibiotic drug) to samples increases the prevalence of *blaCTX-M, blaVIM*, and *Tn916/1545* genes.

It should not be neglected that microorganisms themselves may generate selection pressure regarding antibiotic resistance (Allen et al., 2010). Bacteria from the Actinomycetota produce diverse secondary metabolites, including antibiotics, to promote their survival (Heul et al., 2018). Zhang et al. (2016) observed a strong relationship between the availability of zinc and the presence of *tetQ, tetG, ermB, dfrA,* and *intI1*. Similarly, the presence of bioavailable copper corresponded to the presence of *sul1, ermX ermB, dfrA7* and *aac(6’)-ib-cr*; however, it should be noted that the study was performed in a wastewater treatment plant.

In addition, the abundance of ARGs appears to be contingent on microbial community structure and its shift during processing. While composting is often considered more reliable than storage for reducing pathogen levels in manure, it is also significantly more cost-, time- and labor-consuming (Staley et al., 2021). Zhang et al. (2018) attribute the changes in ARGs levels observed during treatment to the dynamics of the potential host bacteria, with the mobility and type of gene influencing the host range.

The increase in ARGs noted during composting may occur due to the genes being harbored by ARB during proliferation, or HGT between various ARB (Gou et al., 2018). Indeed, genes conferring resistance to chloramphenicol and lincomycin have been found to undergo HGT from gram-positive Actinomycetota to gram-negative Pseudomonadota, despite the clear phylogenetic and ecological boundaries between them (Jiang et al., 2017). HGT is promoted during composting due to it favoring unfavorable conditions for bacterial growth which induce stress-response mechanisms (Lima et al., 2020).

The differences between the ARGs levels detected during composting or storage may be related to the nature of bacterial succession and the survival of ARG-carrying bacteria during the thermophilic phase. During storage, the bacterial strains harboring ARGs were initially suppressed, with their number increasing later on; however, no differences in sample biodiversity were found between treatment strategies. The scale of the study also matters. While lab-scale reactors are simple to prepare and they have clear value in preliminary testing, they nevertheless cannot adequately simulate full-scale processes, and their findings may not directly apply to field-scale studies (Staley et al., 2021).

The detected ARGs could confer resistance to all major antimicrobial classes, such as aminoglycosides, beta-lactams, MLSB, tetracyclines, and vancomycin; all are critical antibiotic groups, with some being even categorized as ‘last-resort’, life-saving antimicrobials. It is important to remember that while the proposed manure management strategies reduce the risk of ARGs spread, they do not eliminate it entirely, and even if not apparent in treated samples, ARGs may persist in the environment below the detection limit. Even trace amounts of genes may further spread with host species succession or be disseminated via HGT.

## 5 Conclusions

Our findings prove that dairy cattle farms can serve as reservoirs of ARGs and MGEs. However, both manure storage and composting, were able to reduce aminoglycoside-, beta-lactam-, tetracycline-, MLSB- and vancomycin-resistance genes as well as MDR genes, MGEs, and various “other” genes; however they had less or no effect on phenicol-, sulfonamide or trimethoprim-resistance genes.

Composting resulted in greater richness and diversity of the bacterial community compared to storage. In addition, while both proposed manure management strategies influence ARGs and MGEs abundance, particular ARGs respond differently to treatment in highly gene-specific ways, and also appear to depend on the dynamics of the bacterial community harboring ARGs. It is difficult to compare the effect of manure treatment strategies due to the range of possible influences. While both strategies reduce ARGs and MGEs levels, composting provides fertilizer with higher biodiversity, and is devoid of pathogens and parasite eggs, but is more labor- and time-consuming than storage. Our findings are important for improving existing or development of appropriate strategies to minimize ARGs dissemination.

## 6 Conflict of Interest

*The authors declare that the research was conducted in the absence of any commercial or financial relationships that could be construed as a potential conflict of interest*.

## 7 Author Contributions

M.Z. - Methodology, Formal analysis, Investigation, Data Curation, Writing - Original Draft, Visualization; A.B. - Formal analysis, Investigation, Writing - Original Draft; M.S. – Investigation, Writing - Original Draft; K.C. - Formal analysis, Data Curation; M.P. - Conceptualization, Writing - Review & Editing, Project administration, Funding acquisition.

## 8 Funding

This work was supported by National Science Centre, Poland (2017/25/Z/NZ7/03026), grant under the European Horizon 2020, in the frame of the JPI-EC-AMR Joint Transnational Call (JPIAMR), JPI-EC-AMR JTC 2017, project INART–“Intervention of antibiotic resistance transfer into the food chain” to MP and partially in the frame of the “Excellence Initiative—Research University (2020– 2026)” Program at the University of Warsaw.

## Acknowledgments

This is a short text to acknowledge the contributions of specific colleagues, institutions, or agencies that aided the efforts of the authors.

## 9 Animal ethics

Following the local legislation and institution requirements, collecting cow manure and conducting research using it does not require consent. The farm owner agreed to the manure being sampled and shared the history of the use of antibiotics on the farm.

## 12 Supplementary Material

Table S1: Humidity [%], pH, and temperature [°C] changes during manure treatments

Table S2. Mean abundance with standard deviation of each phylum across the treatment

Table S3. The results of pairwise PERMANOVA analysis. The multi-testing adjustment is based on Benjamini-Hochberg procedure procedure (FDR)

Figure S1. Core microbiome across analyzed samples

Figure S2. Shannon indexes of microbial community diversity (p-value 0.08; Kruskal-Wallis statistic)

Figure S3. Chao1 indexes of microbial community richness (p-value 0.025; Kruskal-Wallis statistic)

Figure S4. The PCoA of taxon abundance in all sample groups, calculated based on the Bray-Curtis distance; [PERMANOVA] F-value: 7.1007; R-squared: 0.66983; p-value: 0.003

Figure S5. Comparison between resistomes of untreated cattle manure and resistomes at the end of composting and storage

## Data Availability Statement

The datasets generated for this study can be found in the Sequence Read Archive (SRA) repository (NCBI) - BioProject ID: PRJNA1007935.

## Notes

### Competing Interest Statement

The authors have declared no competing interest.

### Summary of Updates

In the new version of the manuscript, the names of the bacteria phyla have been updated.

